# *Title*: Unravelling associations between tree-seedling performance, herbivory, competition, and facilitation in high nature value farmlands

**DOI:** 10.1101/471557

**Authors:** Pedro G. Vaz, Miguel N. Bugalho, José M. Fedriani, Manuela Branco, Xavier Lecomte, Carla Nogueira, Maria C. Caldeira

## Abstract

Herbivory, plant facilitation, and competition have complex impacts on tree regeneration which are seldom investigated together. Grazing exclosure experiments have allowed to quantify the effects of large herbivores on tree regeneration dynamics but have often ignored the effect of herbivorous insects. We experimentally tested how folivory (percentage of leaf damaged by insects), and microenvironment (tree-canopy cover and herbs) jointly alter performance (growth and survival) of seedlings of two dominant Mediterranean oak-species within ungulate exclosures. An agroforestry system dominated by cork oak (*Quercus suber*) and holm oak (*Q. rotundifolia*) was assessed in south-east Portugal. We aimed also to determine whether the two oak-species differ in the interdependences between folivory, microenvironment, covaring factors, and seedling performance. Unexpectedly, under the low–moderate insect defoliation occurred in our 3-year field study, growth and survival of cork and holm oak-seedlings were positively associated with herbivory damage. Herb removal increased oak folivory by 1.4 times. Herb removal was also positively associated with growth, directly and indirectly through its negative effect on oak folivory. Tree-canopy favored insect folivory upon cork oak seedlings directly and upon holm oak indirectly via decreasing light availability. Folivory was threefold greater upon cork than upon holm oak-seedlings. Our study shows that tree-canopy, herbs, and covarying factors can affect cork and holm oak-seedling performances through complex pathways, which markedly differ for the two species. The combined effect of insect herbivory and positive and negative plant-plant interactions need to be integrated into future tree regeneration efforts toward tackling the regeneration crisis of oak agroforestry systems of the Mediterranean.

## 1. Introduction

Tree mortality rates are rising and tree recruitment is increasingly limited (Brasier 1992, Carnicer et al. 2011). Disentangling the associations between the plethora of factors affecting the growth and survival of tree-seedlings is thus becoming an urgent task (Talamo et al. 2015, Gavinet et al. 2016, Kellner and Swihart 2017). Tree-seedling performance is influenced, among other factors, by competition with other plants (Corcket et al. 2003), effects of adult trees (Caldeira et al. 2014), and mammalian and insect herbivory (Leck et al. 2008). Few studies address how insect herbivory, mediated by combined competition and facilitation, affects tree-seedling performance (Sotomayor and Lortie 2015) within mammalian exclosures, pervasively used to protect tree-seedlings (Perez-Ramos and Maranon 2008, Bugalho et al. 2013, Leonardsson et al. 2015, Kellner and Swihart 2017).

Facilitation, such as indirect amelioration of surrounding conditions by adult tree-canopies, and direct competition of herbs, both generate microenvironments affecting tree-seedling performance (Valladares et al. 2016). Relationships between facilitation, competition, and the dynamics of plant communities have long been studied and gained increasing attention (Ladd and Facelli 2005, Callaway 2007, García-Cervigon et al. 2016). Seminal studies have long showed that the probability of tree offspring survival increases with distance from conspecific adults, due to lower predation by host-specific herbivores and pathogens (Janzen–Connell hypothesis; Janzen 1970, Connell 1971). Yet, many facilitation studies focus on effects of light and water on tree-seedling growth and survival (Puerta-Piñero et al, 2007, Rolo et al. 2013), omitting herbivory.

Although herbivory is most often correlated with reduced tree-seedling growth (Marquis 1984, Norghauer and Newbery 2014), there are possible tradeoffs between growth and defense against herbivory (Fine et al. 2006). Plants can maximize their performance by balancing allocation to constitutive defense versus other functions (carbon/nutrient balance hypothesis; Rhoades 1979, Bryant et al. 1983, Bixenmann et al. 2016; but see Hamilton et al. 2001). Shade-tolerant species may allocate resources to defenses following herbivory at the expense of growth rates (resource availability hypothesis; Coley et al. 1985). Plants may also respond to herbivory by over-compensatory growth, increasing growth of aerial parts following herbivory (McNaughton 1983; but see Belsky 1986). Conversely, herbivore species may prefer vigorously growing plants (plant vigor hypothesis; Price 1991) rather than stressed plants (plant stress hypothesis; White 1969, 1974, 1984, Cornelissen et al. 2008). Little is known about the complex effects of insect- and plant-driven conditions on the growth and survival of tree-seedlings inside ungulate exclosures used worldwide to foster woodland restoration.

Cork (*Quercus suber*) and Holm oak (*Q. rotundifolia*) Mediterranean woodlands are high conservation and socio-economic value systems where grazing by domestic or wild mammalian herbivores is very intense. These woodlands extend over 3.1–6.3 million hectares across the Mediterranean Basin (Campos et al. 2013). These multiple-use savanna-like ecosystems are classified under the Pan-European network of protected areas “Natura 2000” (www.natura.org) and considered as high nature value farming systems by the European Environmental Agency (Paracchini et al., 2008). A severe lack of regeneration and adult mortality have been reported in the last decades for cork and holm oak (Brasier 1992, Plieninger et al. 2010, Fedriani et al. 2017). Fenced vertebrate exclosures are a primary practice preventing the major impacts of wild ungulates and livestock (Kellner and Swihart 2017) but very little is known about the impact of folivorous insects on Mediterranean oaks within such exclosures. Furthermore, the relationship between insect damage and seedling chemical defenses (e.g., phenolic compounds) is seldom explored in Mediterranean oaks as well as variation in insect herbivory across seasons and with oak-seedling phenology (Sampaio et al. 2016).

We investigated over three years the complex interplay of insect herbivory and microenvironment inside ungulate exclosures in south-east Portugal toward better understanding the regeneration dynamics of cork and holm oak. We established a two-factor field experimental setting, sowing acorns of the two species beneath and beyond adult tree canopies, where we further manipulated the presence of herbs. The experiment leans on another study investigating the effects of microenvironment on the regeneration of cork and holm oak (Caldeira et al. 2014). We addressed the following questions and associated hypotheses: (i) *Are the performances of cork and holm oak-seedlings (growth, survival) affected by insect herbivory and contrasting microenvironments (tree-canopy, herbs)*? Whereas we expected that insect herbivory generally would diminish growth and survival of seedlings (Stephens and Westoby 2015, Tiberi et al 2016), we predicted that performance would vary among facilitation and competition conditions created by adult tree-canopies and herbs, respectively (Caldeira 2014). (ii) *Does herbivory vary intra-annually, with microenvironment, and in response to seedling chemical defenses?* Herbivory would be greater in spring, following seedling leaf development phenology (Aide 1993, Lamarre et al. 2014, Tiberi et al. 2016), would vary among microenvironments as these affect insect feeding activity (Valladares et al. 2016), and most damaged seedlings would have higher levels of phenols (Feeny 1970, Bryant et al. 1987). (iii) *Are there differences between oak-species in the interdependence between herbivory, tree-canopy, herbs, covarying factors (light, litter biomass, oak resprouting status), and seedling performances?* Because we are addressing two coexisting and closely related oak-species (Pearse and Hipp 2009), we would expect similar responses to insect herbivory and a similar network of direct and indirect associations affecting seedling performances.

## 2 Materials and methods

### 2.1 Field methods

#### 2.1.1 Study area

We conducted this study at Tapada Real de Vila Viçosa (38°47’N, 7°22’W), a 900-ha estate in south-east Portugal. The local climate is Mediterranean with hot, dry summers and cool, wet winters. Annual precipitations during the study period (November 2003 to October 2006) were 639, 469, 442, and 739 mm falling mainly in fall and winter, while mean annual temperature is 16.0 °C, ranging from 8.3 °C in December–January to 24.0 °C in July–August. Soils are poorly developed haplic leptosols (WRB 2006) with dominant bedrock of schist (Caldeira et al. 2014). The experimental plots were established in homogeneous fenced areas of a mixed cork oak and holm oak woodland. There were 3–4 cork oak and 3–5 holm oak trees in each fenced area. Cork and holm oak are slow-growing, shade-tolerant species. They do not grow in winter, have a growth flush of new leaves in spring (synchronized with leaf shedding from the previous year), grow slowly or stop growing in summer, and have a second but less intense growth flush in fall. New leaves last ~ 12 to 25 months (Díaz et al. 2004, Pinto et al. 2011). Grasslands were mainly composed of annual species and the shrub understory was dominated by rockrose (*Cistus ladanifer*).

#### 2.1.2 Experimental design

We pre-germinated cork and holm oak acorns in a greenhouse until the radicles were 0.5 cm long. These acorns were collected in the study area from beneath at least 5 different cork and holm oak trees. Cork and holm oak acorns weighed, on average, 3.5 g ± 0.9 (fresh mass range) and 2.8 g ± 0.8, respectively. We buried the acorns 2 cm deep into the soil 30 cm apart in field experimental plots. The sowing was done within typical exclosures that prevent access from large herbivores (red deer, *Cervus elaphus*; fallow deer, *Dama dama*; and wild boar *Sus scrofa*) and thus facilitate tree regeneration. Specifically, we held the field experiment within five 25×25 m fenced areas (2.2 m height) established in previous studies (e.g., Bugalho et al. 2011, Caldeira et al. 2014). To assign a tree-canopy treatment (levels: beneath, beyond), we haphazardly established 18 plots (2×4 m), 10 beneath the canopy of mature *Quercus* trees and eight beyond trees in open grassland areas. Owing to space constraints, the number of plots was not the same in all fences. Four plots were established in two fences, while the remaining fences had five, three, and two plots each. These fenced areas were established in homogeneous areas with similar slopes (4.6° ± 0.5° SE) at NW-SE orientations. Distance between fenced areas was 250-400 m (Lecomte et al. 2016). To establish an herb presence treatment (present, removed), each plot was halved into two subplots (2×2 m), in one of which the herb vegetation was removed by hand thrice a year. In mid-November 2003, we sowed 18 acorns of each species per subplot interspersing pregerminated acorns of the two species. The total 1296 planted seedlings were tagged and monitored thereafter.

#### 2.1.3 Data collection

In a first phase of the experiment, each seedling was monitored 18 times, nearly monthly from April 2004 to June 2005, and then every 2–4 months onwards until July 2006. Seedling mortality was recorded when we observed the senescence of all leaves. As for growth, because these oak species often experience above-ground mortality of all tissues followed by resprouting, we examined the longest period of seedling height growth (hereafter growth) uninterrupted by the complete loss of seedling aerial parts. For analysis, we derived growth (mm/day) from the quotient of the difference between final and initial plant total height (main stem length) by the number of days. Over that time period, at each monitoring date, we recorded the number and the percentage of leaves having >25% area of damage attributable to insect herbivores (chewed, mined, or skeletonized leaves and galls). We disregarded chlorotic and necrotic spots that may suggest attack by pathogens or micro-organisms. To assess overall seedling size, we recorded its total number of branches coming off the main stem at each date. For analysis, we derived per seedling the mean percentage of leaves damaged by insects and the mean number of branches. To assess the potential resprouting effect on seedling performance and herbivory (Fornoni 2011), we recorded resprouting status during the experiment as binary (yes/no). Seedlings that did not resprout since 384 days by the end of the experiment were assumed to be dead. We opted for this cutoff for analysis as only 5% of observed resprouting events occurred after 384 days.

To evaluate whether leaf litter layer and light were related to seedling performance and herbivory (Fine et al. 2006, Valladares et al. 2016), we proceeded as follows. We collected litter samples (senesced tree leaves) in 2004 from a random 10×50 cm area in all subplots, which were oven-dried at 70 °C until attaining constant weights. We measured light availability (proportion of light reaching seedlings) using hemispherical photographs taken with a fish-eye lens and a digital camera. We took photographs at dawn in the summer 2004, ~60–70 cm above the herb layer at the midpoint of each subplot. Images were analyzed with Hemiview software (Delta-T Devices Ltd, Cambridge, UK), and the proportion of global (direct and diffuse) radiation beneath tree-canopy relative to the open (global site factor, GSF) was determined. Additional details are provided in Caldeira et al. (2014).

In the second phase of the experiment (i.e., during 2006), we subsampled 222 seedlings from the original plantings, on which we assessed insect herbivory in May (spring measurement), July (summer), and October (fall). The 222 seedlings were distributed randomly across all field plots. The subset was balanced by tree-canopy, herbs, and species treatments. To assess the relationship between insect damage and seedling chemical defenses, we measured the concentration of phenolic compounds on the 222-seedling subset. Two fresh intact mature leaves per seedling were collected and immediately frozen in liquid nitrogen. Total phenols concentration was determined based on the Folin-Ciocalteau method (Folin and Ciocalteu 1927) using 20 mg of material. Callibration curves using standard gallic acid solutions were created. Total phenols were expressed in mg of gallic acid equivalents per dry weight.

To evaluate herbivory on the 222-seedling subset, one branch per plant was tagged and monitored. On the three dates, we recorded the total number of leaves at the branch and the estimated percentage of damage attributable to insect herbivores on each leaf. For analysis, we calculated an index of herbivory for the branch at each date (Chacón and Armesto 2006). Each leaf was assigned to a category of damage according to the leaf area affected: 0=0%, 1=1–15%, 2=15–50% and 3= 50–100%. The herbivory index (HI) was defined as *HI* = Σ*C_i_*×*n_i_*/*N*, where *C_i_* corresponds to the midpoint of each damage category (i.e., *C_1_* =8, *C_2_* = 33, and *C_3_*=75%, respectively), *n_i_* is the number of leaves in the *i*^th^ category of damage, and *N* is the total number of leaves in the branch.

### 2.2 Statistical analyses

#### 2.2.1 Effects of herbivory and microenvironment on oak-seedlings

##### Growth

To evaluate the effects of insect herbivory on growth of oak-seedlings as affected by tree-canopy, herbs, and oak-species, we used linear mixed-effects modeling (LMM). We also tested the effects on growth of the covariates — mean number of branches, resprouting status, biomass of litter, and light availability. Because the sampling design was intended to capture the canopy × herbs × species interaction, we included this interaction in the LMM. Prior to analyses, we used the equation 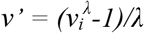, the Box-Cox family of transformations on each *i*^th^ value to find the best transformation (Quinn and Keough 2002) on growth (*λ* = 0.30) meeting the assumption of normally distributed errors. Each variable was expressed per emerged oak. A matrix of Spearman’s correlations for initial explanatory variables revealed litter was correlated with light (*r_s_* = −0.78, p < 0.001) and was excluded from this analysis to prevent collinearity. Spatial groupings of plots were nested within fences as experimental units, so we used both as random factors in LMM following that hierarchical order. However, a likelihood ratio test (*anova* command in *nlme* package in R; Pinheiro et al. 2017) suggested adding fence to random effects was not a significant improvement (*L* = 4.88, *P* = 0.068; Zuur et al. 2009). Furthermore, we tested if the linear regression model would be preferable to the LMM with plot as random factor (*exactLRT* command, *RLRsim*; Scheipl et al. 2008) and opted for the latter (*L* = 14.28, *p*<0.001). We used identity link to fit LMM. The minimal adequate (optimal) LMM was arrived at by first fitting the full model (with all the abovementioned explanatory terms simultaneously) followed by backward elimination of non-significant (*P* > 0.05) explanatory variables one at a time and then applying the likelihood ratio of nested models (Zuur et al., 2009). We evaluated model adequacy by checking normal distribution of residuals, plotting residuals vs fitted values and explanatory variables and we evaluated model fit by the marginal *R*^2^ (proportion of variance explained by the fixed effects; Nakagawa and Schielzeth, 2013).

##### Survival

To identify multivariate predictors for the relative risk of death of cork oak and holm oak-seedlings, we used Cox proportional hazards frailty modeling (Therneau and Grambsch 2000). The response variable was the number of follow-up days until seedling death and was modeled as right-censored due to the uncertainty that seedlings could eventually die after the experiment. We used the same full set of predictors as in the full LMM for growth. The effect of experimental plot was accounted for by including it into the model as a random proportionality factor (“frailty”). To fit the Cox regression, we used the *survival* package in R (Therneau 2015). The significance of each predictor and interaction was evaluated by backward-stepwise elimination from the full model (Therneau and Grambsch 2000). Because light availability was the only non-significant (chi-square test; *P* > 0.05) explanatory variable, we assessed the final model by dropping light. Also, resprouting status was removed from the model owing to a strong violation of the Cox proportional hazards assumption assessed by smoothed scaled Schoenfeld residual plots, even after examining an interaction between resprouting and time (Therneau et al. 2016). Yet, we examined below the effect of resprouting on seedling death (see Path analyses).

#### 2.2.2 Predictors of herbivory

To assess how herbivory index (HI) changed over time with the explanatory variables canopy × herbs × species interaction, light availability, and phenols, we used the 222-seedling subset to construct a LMM with identity link. Litter was again omitted to prevent collinearity. To test for differences of herb and canopy effects among seasons, we included herbs × month and canopy × month interactions into the full suite of fixed explanatory terms. Since HI was derived thrice in one branch per plant (in May, July, and October), seedling was incorporated into the random part of the model nested within experimental plots (plot / seedling). The response variable HI was logit-transformed prior to analyses (Warton and Hui 2011). Procedures for model fitting and adequacy inspecting were similar to the above-mentioned LMM for seedling growth.

#### 2.2.3 Path analyses

We tested for direct, indirect, or cascading effects among variables by performing a confirmatory path analysis (*sensu* Grace 2006) using the D-step procedure. A path diagram denotes by arrows which variables are influencing (and influenced by) other variables. This procedure tests whether any paths are missing from the model, and whether it would be improved with the inclusion of any missing path (Shipley, 2009). We included the mean number of branches, biomass of litter, resprouting status, and light availability as explanatory variables. Phenols were not included in this analysis since they were measured in the 222-seedling subset but not in the whole dataset used here. Likewise, herbivory was assessed by the percentage of leaves with >25% area damaged and not by HI. To assess how different between oak-species were the pathways of the dependencies among variables, we constructed one model per oak-species.

For both models, we first expressed the hypothesized relationship between variables in the form of a path diagram. Based on our previous knowledge of the system (e.g., Bugalho et al. 2011, Caldeira et al. 2014), we hypothesized the following direct relationships: (i) herbivory affecting (negatively, −) growth and (positively, +) death events; (ii) herbs, canopy, and number of branches affecting seedlings’ growth (−, +, +, respectively), death (+, −, −), and insect herbivory (−, +, +); (iii) litter biomass affecting (+ or −) herbivory; (iv) resprouting affecting (−) death; (v) canopy affecting (−) light availability. Indirect effects would include canopy and herbs both affecting herbivory by affecting (+) litter biomass.

We treated tree-canopy, herbs, death, and resprouting as binary variables and centered and standardized the mean number of branches, litter, and light. Using the R package *piecewiseSEM* (Lefcheck, 2016), the path diagram was translated to a set of six mixed-effects models explaining growth, herbivory, death, mean number of branches, litter biomass, and light availability. We entered plot as the random factor in all six models. Growth (Box-Cox-transformed; *λ* = 0.30), mean number of branches, litter biomass, and light availability were fitted by three LMM with identity links (*nlme* package in R; Pinheiro et al., 2017). Death and percentage of damaged leaves were both fitted using binomial generalized LMM (GLMM) with logit links (*lme4* package; Bates et al., 2015). In the latter case, we used the seedling total number of leaves as a weight argument. We tested for the validity of the hypothesized relationship pathways using the C-statistic (Shipley, 2009). Once the generalized path model was validated, we obtained the path coefficients by fitting the models. We did all analyses in R v. 3.3.3 (R Core Team, 2017).

## 3 Results

Almost 80 % of sowed acorns (1019 out of 1296) had emerged by the end of the study period, ~ 60% of which survived (cork oak = 335 of 578; holm oak = 260 of 441). Seedlings grew at 0.032 mm/day (± 0.0001 SE), reaching a mean height of 14.7 cm (± 0.2 SE; cork oak = 15.3 cm ± 0.3, holm oak = 14.0 cm ± 0.3). Mean insect defoliation during the experiment was 7.3 % of seedling leaf area (± 0.3 SE; cork oak = 10.9 % ± 0.6, holm oak = 3.8 % ± 0.3).

### 3.1 Effects of herbivory on growth and survival

#### 3.1.1 Growth

When we controlled for the effect of random factor (i.e., experimental plot), our optimal mixed model (Table 1) showed that greater growth rates were significantly associated with greater herbivory in both oak-species. For example, seedling growth with 45% damaged leaves exceeded by twofold that of seedlings with 10% damaged leaves (Fig. 1A). Also, growth rates were significantly higher in larger seedlings – e.g., it doubled from three- to six-branch seedlings (Fig. 1B). In addition, the effect of herbs was significant (*F*_1, 1006_ = 25.80, *P* < 0.001; Type-III *F*-test, Satterthwaite approximation for denominator degrees of freedom), with herb removal increasing growth by 1.9 times (Fig. 1C). The effect of the canopy × species interaction was significant (*F*_1, 1005_ = 4.75, *P* = 0.029), i.e., the effect of tree-canopy was not the same for both species. Beneath tree canopies, holm oak growth ratio was 2.2 fold greater than that of cork oak (Fig. 1D). No other variable or interaction was found to be significant for seedling growth.

**Figure 1.**
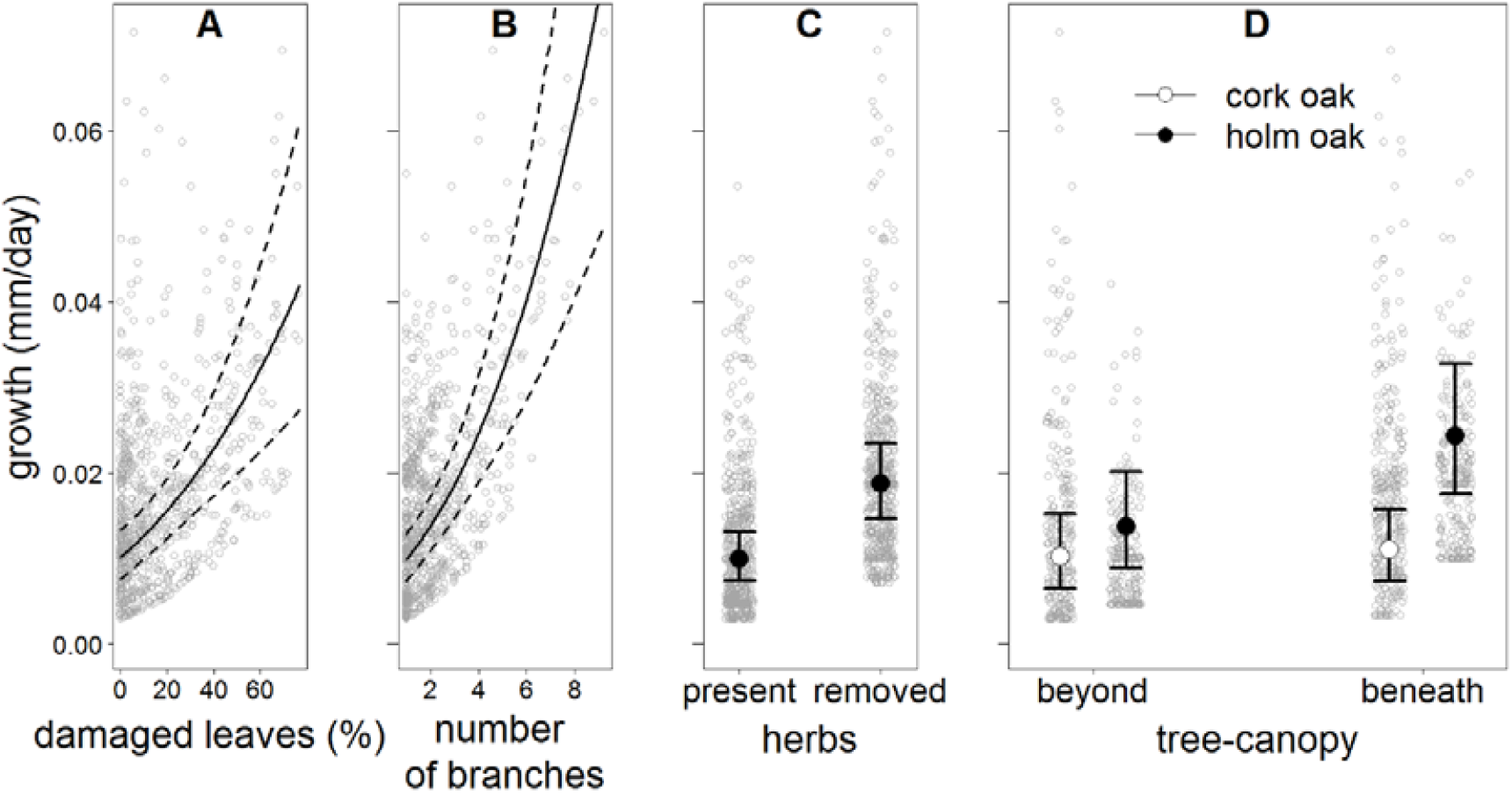
Mean (± 95% CI) fitted values for the significant terms of the optimal mixed-effects model predicting cork oak and holm oak-seedlings growth. Grey dots are predicted values.

**Table 1.** Fixed part of the optimal mixed-effects model predicting cork oak and holm oak-seedlings’ growth rate. SE = standard error; CI = effect size confidence intervals.

| Variable | marginal $R^2 = 0.11$, conditional $R^2 = 0.15$ | | | | | |
| --- | --- | --- | --- | --- | --- | --- |
| | Estimate | SE | $t$ | 2.5% CI | 97.5% CI | $P$ |
| intercept | -2.819 | 0.066 | -42.61 | -2.946 | -2.692 | <0.0001 |
| herbs (removed) | 0.172 | 0.034 | 5.08 | 0.106 | 0.238 | <0.0001 |
| tree-canopy (beneath) | 0.019 | 0.074 | 0.25 | -0.124 | 0.162 | 0.800 |
| species (holm oak) | 0.079 | 0.052 | 1.52 | -0.024 | 0.179 | 0.129 |
| percentage of damaged leaves | 0.006 | 0.001 | 5.03 | 0.003 | 0.008 | <0.0001 |
| number of branches | 0.087 | 0.014 | 6.36 | 0.060 | 0.113 | <0.0001 |
| canopy (beneath) : species (holm oak) | 0.150 | 0.069 | 2.18 | 0.016 | 0.286 | 0.029 |

An additional exploration of raw data showed a weak correlation (Spearman rank correlation, *r_s_* = 0.19, *P* < 0.001, *n* = 1019) between percentage of herbivory and seedling size (number of branches). Also, a 3D visualization of growth, herbivory, and seedling size showed no obvious pattern, except that holm oak seedlings had less herbivory damage and fewer branches (Fig. A.1, Appendix A).

#### 3.1.2 Survival

Cox regression analysis (Table 2) indicated that seedling survival was significantly positively associated with insect herbivory. Every 10% increment in damaged leaves was associated with a 40% lower hazard ratio (Fig. 2A). The analysis likewise showed that the effect of seedling size was significant, with an additional seedling branch being associated with a 50% lower hazard ratio (Fig. 2B). The main effect of herbs was significant as well (*χ^2^* = 20.61, df = 1, *P* < 0.001 0.001; Type-III chi-square test), with seedlings growing in bare soil having 40% lower hazard ratios than amid herbs (Fig. 2C). Also, herbs significantly interacted with both tree-canopy and seedling species effects (*χ^2^* = 76.74, df = 12, *P* < 0.001). Specifically, with herbs present beyond canopies, death was 35% less likely to occur in cork than in holm oak. With herb removal beyond canopies, death was 34% less likely to occur in cork than in holm oak. With herb removal beneath canopies, death was 27% less likely to occur in holm than in cork oak-seedlings (Fig. 2D). No other variable was found to have a significant effect on seedling survival.

**Figure 2.**
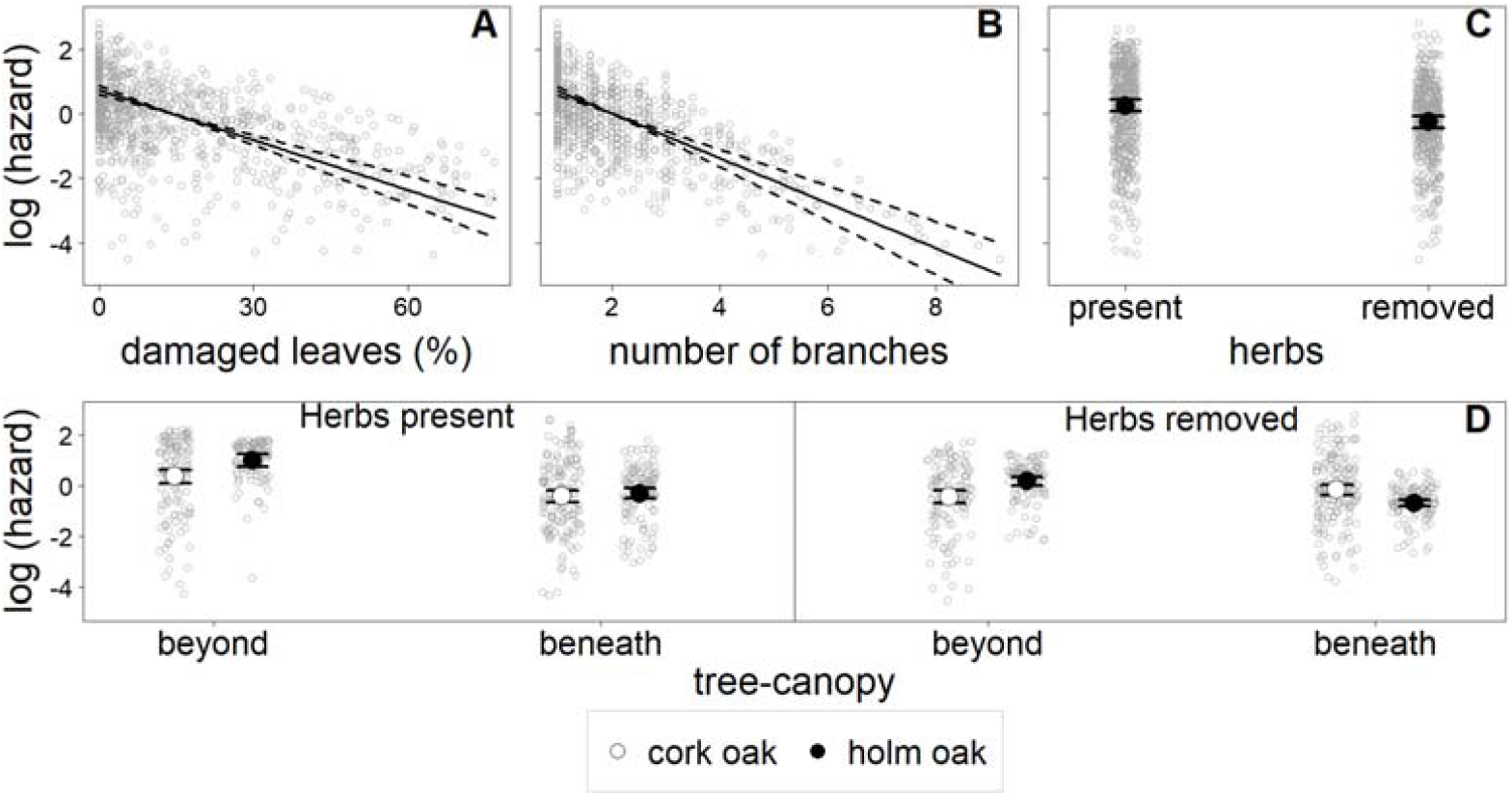
Mean (± 95% CI) fitted values for the significant terms of the optimal Cox regression model predicting the (logarithm of) mortality rate for cork oak and holm oak-seedlings. Grey dots are predicted values.

**Table 2.** Optimal survival model by Cox multiple regression for relative risk of death of cork oak and holm oak-seedlings. Plot was used as a random proportionality factor (frailty). HR = hazard rate.

| Variable | Likelihood ratio test = 431.9, $P < 0.0001$ | | | | |
| --- | --- | --- | --- | --- | --- |
| | β-coef. | SE | $\chi^2$ | $P$ | HR (95%CI) |
| tree-canopy (beneath) | -0.095 | 0.299 | 0.10 | 0.751 | 0.90 (0.51–1.63) |
| herbs (removed) | -0.506 | 0.189 | 7.21 | 0.007 | 0.60 (0.42–0.87) |
| species (holm oak) | -0.616 | 0.164 | 1.48 | 0.111 | 0.54 (0.39–1.75) |
| percentage of damaged leaves | -0.052 | 0.005 | 109.25 | <0.0001 | 0.95 (0.94–0.96) |
| number of branches | -0.694 | 0.070 | 97.98 | <0.0001 | 0.50 (0.44–0.57) |
| tree-canopy (beneath) : herbs (removed) | 0.704 | 0.273 | 6.67 | 0.0101 | 2.02 (1.18–3.45) |
| tree-canopy (beneath) : species (holm oak) | -0.205 | 0.272 | 0.57 | 0.451 | 0.81 (0.48–1.34) |
| herbs (removed) : species (holm oak) | -0.065 | 0.275 | 0.06 | 0.812 | 0.94 (0.55–1.61) |
| canopy (beneath) : herbs (removed) : species (holm oak) | -0.832 | 0.417 | 3.98 | 0.036 | 0.44 (0.19–0.99) |
| frailty |  |  | 47.68 | <0.0001 |  |

### 3.2 Predictors of herbivory

Our LMM for insect herbivory (Table 3) revealed significant differences related to seedling species (*F*_1, 173_ = 77.47, *P* < 0.001, Type-II *F*-test). Specifically, the herbivory index (HI) was 2.8 fold greater in cork than in holm oak-seedlings (Fig. 3A). Likewise, the effect of herbs was significant (*F*_1, 173_ = 11.19, *P* = 0.001), with herb removal significantly increasing HI by 1.4 times (Fig. 3B). Herbivory also significantly changed over seasons (*F*_2, 382_ = 49.18, *P* < 0.001), decreasing 21% in July as compared to May, and peaking in October, when it was 1.3 and 1.7 greater than in May and July, respectively (Fig. 3C). Our LMM likewise showed that light significantly affected herbivory. For example, HI decreased by 12% from seedlings with 0.2 to seedlings with 0.4 proportion of light available (Fig. 3D). Leaf phenol content was not deemed a significant predictor of HI.

**Figure 3.**
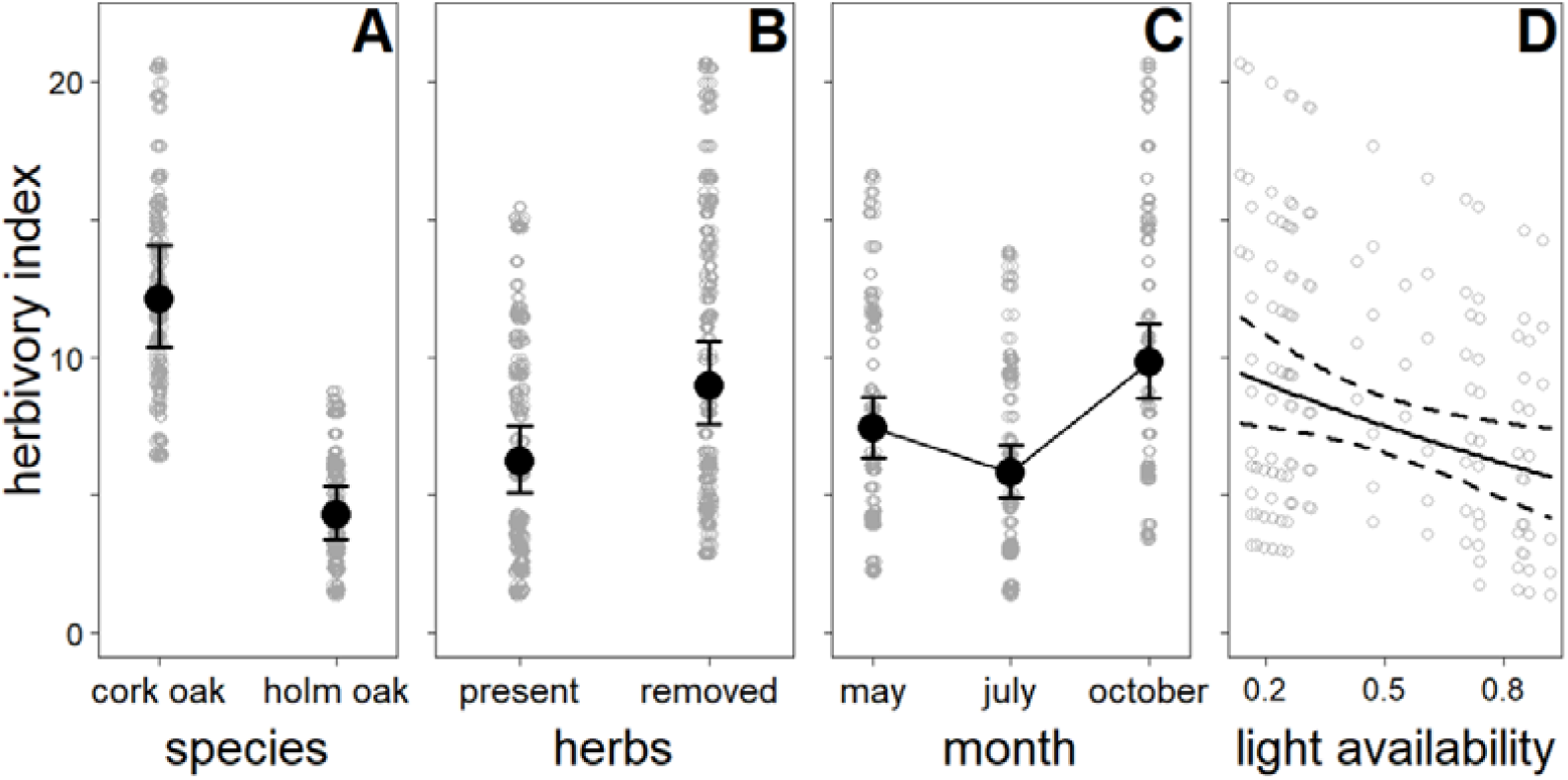
Mean (± 95% CI) fitted values for the significant terms of the optimal mixed-effects model predicting index of insect herbivory on cork oak and holm oak-seedlings in May, July, and October. Grey dots are predicted values.

**Table 3.** Fixed part of the optimal mixed-effects model predicting the index of insect herbivory on cork oak and holm oak-seedlings in May, July, and October.

| Variable | marginal $R^2 = 0.32$, conditional $R^2 = 0.77$ | | | | | |
| --- | --- | --- | --- | --- | --- | --- |
| | Estimate | SE | $t$ | 2.5% CI | 97.5% CI | $P$ |
| intercept | -1.724 | 0.145 | -11.88 | -2.005 | -1.443 | <0.0001 |
| species (holm oak) | -0.839 | 0.095 | -8.80 | -1.024 | -0.653 | <0.0001 |
| month (July) | -0.189 | 0.043 | -4.40 | -0.273 | -0.105 | <0.0001 |
| month (October) | 0.240 | 0.043 | 5.54 | 0.155 | 0.324 | <0.0001 |
| herbs (removed) | 0.300 | 0.098 | 3.05 | 0.110 | 0.489 | 0.003 |
| proportion of light available | -0.526 | 0.209 | -2.52 | -0.949 | -0.120 | 0.012 |

### 3.3 Network of direct and indirect effects

Path analyses confirmed a direct and positive association between herbivory and seedling growth in both oak species (Fig. 4). Herbivory association with seedling death was significantly higher in cork oak, conveying the largest effect (−7.40) between variables (Fig. 4A). That path was not deemed significant in holm oak (*P*>0.05) and so it was dropped (Fig. 4B). These analyses confirmed the significance of herb removal on increasing growth and herbivory in both species (Fig. 4A-B), while decreasing death events in cork oak only (Fig. 4A). Tree-canopy affected positively herbivory upon cork oak (Fig. 4A), whereas it affected growth positively and death events negatively in holm oak (Fig. 4B). There were also indirect and significant effects of tree-canopy on holm oak herbivory through reduction of light availability (Fig. 4B), whereas this indirect effect was not observed in cork oak (Fig. 4A). Noticeably, seedling resprout was positively associated with cork oak herbivory but negatively associated with holm oak herbivory (Fig. 4A and 4B). Seedlings with higher growth rates were also less likely to die in both species (Fig. 4A–B).

**Figure 4.**
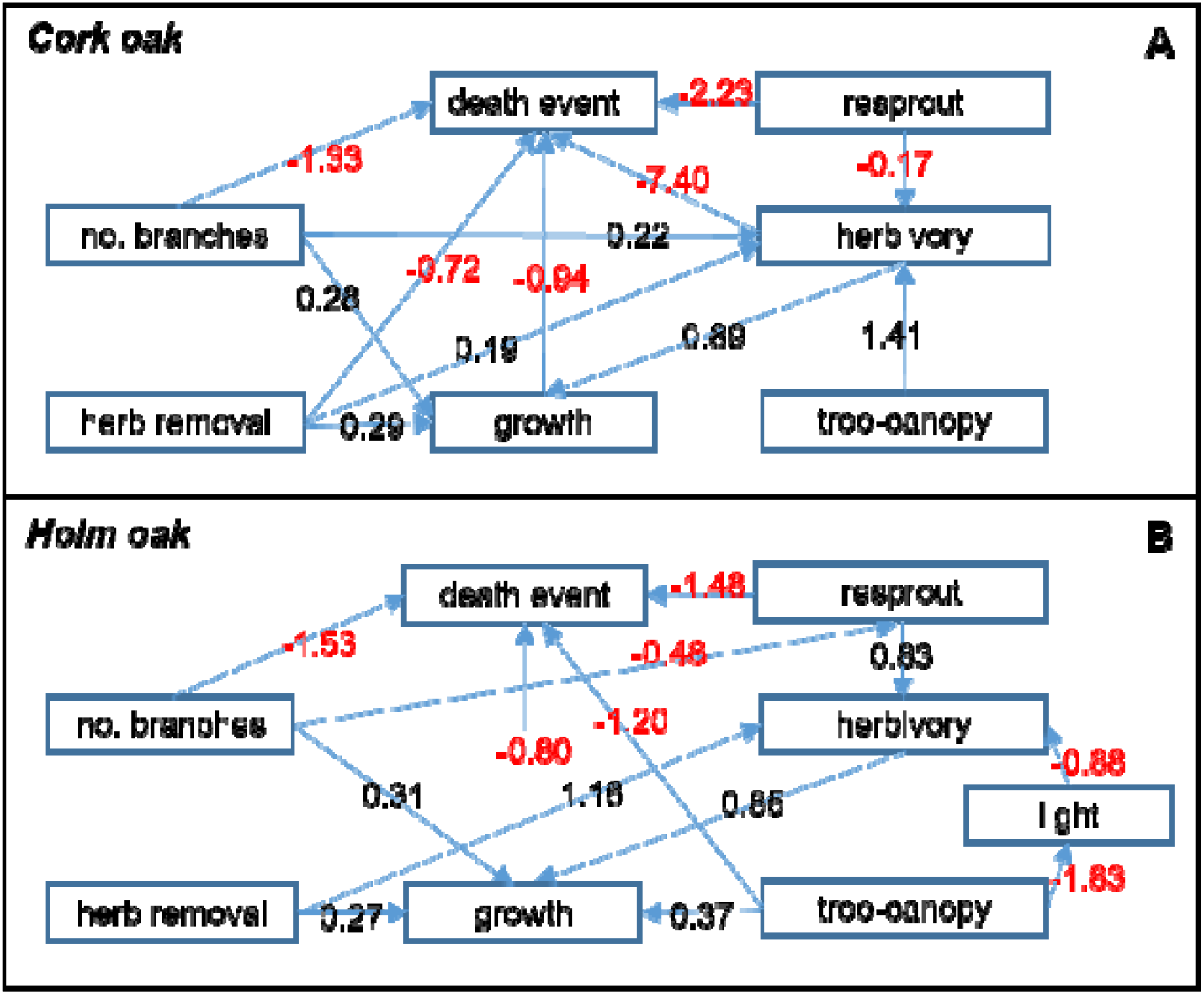
Final path diagram for cork oak (*P* = 0.697, n = 578; A), and for holm oak (*P* = 0.232, n = 441; B) seedlings. Arrows represent directed relationships between variables (paths) significant at the 0.05 level, each having a standardized coefficient (sign indicates whether the relationship is positive or negative for that direct effect). Coefficients can be compared to assess the relative effects of the variables.

## 4. Discussion

### 4.1 Herbivory direct responses

Overall, the levels of insect defoliation during the experiment were low to moderate (Crawley 1997), although they were variable between species. Our prediction of intra-annual herbivory variation was confirmed by the analyses. Indeed, findings describing patterns of herbivory observed in May and July agree with previous research stating that most cork and holm oak folivores feed on expanding leaves during spring and to a lesser extent in summer (Tiberi et al. 2016). Feeny (1970), also stressed that most defoliators prefer feeding on young rather than older leaves. Although late-season defoliators are unusual in our system, plant growth may occur after first rainfalls, which supports the peak of herbivory observed in October in our study.

Herb removal enhanced folivory in our site through several possible mechanisms. First, generalists may have aggregated on seedlings in the absence of herbs nearby. Second, host location by insect specialists may have been easier also on isolated seedlings rather than mixed amid herbs, both visually and chemically. Previous research has shown that specialists can be particularly sensitive to host location and conspicuousness (Bertheau et al. 2010). Lastly, neighbor herbs in the study area may accommodate several predators, such as Syrphidae larvae and Coccinellidae adults, which we suggest may have reduced the number and activity of oak folivores (see review by Schmitz 2010). Broadly, oak-seedling-herb vegetation associations in our system are consistent with the widespread associational resistance model (Bertheau et al. 2010).

Response of herbivory to tree-canopy was oak species-specific. Only for cork oak it directly increased folivory as initially hypothesized, having had an indirect impact on holm oak herbivory via decreasing light availability. For cork oak, we suggest conditions beneath tree-canopy other than less light were likely dictating folivory. Cork oak insect specialists may have been more common near adult trees, corroborating the Janzen-Connell hypothesis (Janzen 1970, Connell 1971). In contrast, factors co-varying with light (e.g., temperature, humidity, olfactory or visual environment) were likely playing a role on patterns of folivory upon holm oak, as previously shown by Sipura and Tahvanainen (2000) for willow trees. Others have also demonstrated light affects the abundance and performance of herbivores, both indirectly through effects on host plants (Barber and Marquis 2011) and directly (Martinat 1987, Mazza et al. 2002). Our holm oak results adhere to the widely observed pattern that shadier conditions increase susceptibility of plants to insect herbivory (Valladares et al. 2016). On the other hand, it is well established that plants growing under lower light contain relatively more nitrogen and less carbon-based secondary compounds such as phenols (Dudt and Shure 1994), and are therefore better food for insects (Sipura and Tahvanainen 2000), as also substantiated in the theory of plant resource allocation (Bryant et al. 1983, Herms and Mattson 1992). However, our outcome may not fit this rational as we did not detect any measurable effect of phenols on folivory, though these compounds are among the most general chemical barriers of woody plants against herbivores (Feeny 1970, Bryant et al. 1987). Still, it remains possible that we may not have detected differences in phenols in the undamaged leaves used for sampling, as defenses could have been induced in response to herbivory damage only. More work is needed to assess the mechanisms through which light environment drive folivory, including leaf nutritional and mechanical properties (Onoda et al. 2011).

The two oak species showed a noticeable difference in folivory levels. Although plant defense usually relies on multiple traits (Agrawal and Fishbein 2006), stronger damage upon cork oak leaves in our results may have been partly due to its softer leaves relative to holm oak (Caldeira et al., unpublished data). Also, divergence in timing of leaf growth between co-occurring plants warrants future consideration in terms of its implications for differential folivory. In the study area, yearly holm oak budburst can occur six weeks earlier than cork oak (Pinto et al. 2011). Herbivores are more active in late spring, so new cork oak leaves may have been preferred over developed and hardened holm oak leaves. Moreover, herbivory on both species had opposite responses to resprout. For cork oak, the negative effect of resprout on folivory suggests resprouting is worth investigating as one possible mechanism of herbivory tolerance (Fornoni 2011). Overall, we provided evidence that these two closely related species (Pearse and Hipp 2009) may diverge in their relationships between early performance and insect herbivory.

### 4.2 Herbivore-mediated effects on seedling performance

Surprisingly, oak-seedling growth and survival were positively associated to insect herbivory. Oak-seedlings seemed to have been capable of compensating for herbivory tissue loss through increased growth. This challenges the common assumption that plants growing in low resource environments, like in our study area, are worst at compensating for or tolerating herbivory (Bryant et al. 1983, Coley et al. 1985, Crawley 1997). However, because the level of insect defoliation was low to moderate, extrapolating our results to situations of severe herbivory must be considered cautiously (but see Díaz et al. 2004). Another possibility is that folivores consumed seedlings in better physiological condition having higher nutritional value, which were also more likely to have higher growth rates. Since these folivores feed mostly on expanding leaves, they may have preferred seedlings growing vigorously (Price 1991, White 2009). Yet, we found no obvious pattern in a three-way plot of growth rate, herbivory, and seedling size (Fig. A.1, Appendix A).

Under the low–moderate folivory in this study, we found herb removal was consistently positively associated with growth directly and indirectly through increasing herbivory in both oak species. This result provides useful information on how plant regeneration may be modified through consumer-mediated effects, a less-explored topic (Chaneton et al. 2010). Additionally, this lends contribution to the substantial work suggesting neighboring vegetation has potential to change establishment dynamics by affecting plant growth (e.g., Cuesta et al. 2010, Caldeira et al. 2014). However, path analyses showed the indirect effect of herb removal on seedling growth via increasing herbivory could have exceeded its positive direct effects. Thus, higher growth rate in subplots with no herbaceous cover, rather than representing just a release from competition with annual herbaceous, may have been chiefly explained as an indirect effect of herbs on decreasing the impact of insect herbivores on focal oaks. This was an asset of our results, as indirect effects commonly lead to a misinterpretation of experimental data on direct effects (see Burger and Louda 1994).

The 40% lower hazard ratio (likelihood of death) for seedlings on subplots with no herbaceous cover in our results corroborates Caldeira et al. (2014) who, despite not analyzing folivory effects, have demonstrated that herb biomass can negatively affect oak-seedling survival. In turn, effects of canopy-tree on survival, jointly interacting with herbs and species, and adding to the effect of folivory, exposed the complexity of factors influencing the survival of these oak-seedling species. An overall higher hazard ratio for holm oak seedlings beyond tree-canopies is in line with Rodriguez-Calcerrada et al. (2010) results for *Quercus pyrenaica* and *Q. petraea*, where lowest seedling survival occurred beneath closed tree-canopies. But our models showed clearly that the mechanistic pathways of tree-canopy microhabitat on seedling growth and survival via folivory varied for both oak species. In cork oak, the large effect of tree-canopy fostering folivory indirectly promoted seedling performance through strong associations of folivory with both raising seedling growth and survival. Conversely, in holm oak, tree-canopy microhabitat enhanced markedly seedling performance by a simple direct and an indirect cascade mechanism, both enhancing seedling establishment. Through the latter, tree-canopy decreased light, which reduced herbivory, which, in turn, associated with higher holm oak growth rates. Holm oak seedlings are more shade-tolerant than cork oak (Petroselli et al. 2013), likely explaining higher holm oak growth rates beneath tree canopies in our results. Broadly, our results are relevant in determining the regeneration dynamics of these investigated Mediterranean oaks, further adding contribution to the flourishing interest in indirect ecological interactions (Sotomayor and Lortie 2015).

## 5. Conclusions

Insect herbivory does not appear to negatively affect growth and survival of cork and holm oak-seedlings at low–moderate levels registered in our site. On the contrary, cork and holm oak performance (survival and growth rate) can be positively associated with insect herbivory, which can be amplified beneath adult oak trees and especially by herb removal. From an applied and woodland restoration perspective, this implies that tree-seedling performance inside ungulate exclosures can be optimized by reducing herb cover (e.g., mowing) and by establishing oak-seedlings near adult tree-canopy microenvironment. Circumventing the debate on the use of insecticides to reduce herbivory in forests, our study suggests alternative management practices may be available. Further research evaluating whether and how the associations found in this study change under greater levels of insect herbivory is needed. Plus, our findings support that there is intra-annual herbivory variation upon these dominant Mediterranean oak-seedling species. Periods of great leaf expansion and greater folivory appear to be synchronous. Regrettably, forecasted Mediterranean climate change scenarios of increasing frequency of high temperatures and severe droughts will likely affect oak regeneration niches as well as seasonal rates of oak folivory, with unclear consequences for competition and facilitation interactions. Lastly, our study highlights that herbivory, tree-canopy, herbs, and covarying factors can affect cork and holm oak-seedling performances through complex pathways, which markedly differ for the two species. The joint effect of insect herbivory and positive and negative plant-plant interactions thus need to be integrated differently for the two species into future tree regeneration efforts using ungulate exclosures. Broadly, plant facilitation (e.g., tree-canopy and nurse effects) and competitive interactions need to be carefully considered alongside herbivory when tackling the Mediterranean oak regeneration crisis.

## Authors’ contributions

Conceived the study and designed the experiment: MCC MNB MB. Performed the experiment: XL CN MCC. Analyzed the data: PGV. Wrote the paper: PGV MNB JMF MCC MB XL CN.

## Acknowledgements

We thank Fundação da Casa de Bragança for granting access to Tapada Real de Vila Viçosa. We appreciate critical comments by Dr. Markus Eichhorn, University of Nottingham, and other five anonymous reviewers, which made numerous useful comments that improved previous drafts.

## References

Agrawal AA, Fishbein M (2006) Plant defense syndromes. Ecology, 87, S132-149.

Aide TM (1993) Patterns of leaf development and herbivory in a tropical understory community. Ecology, 74, 455-466.

Bale JS, Masters GJ, Hodkinson ID, Awmack C, Bezemer TM, Brown VK, … Whittaker JB (2002) Herbivory in global climate change research: direct effects of rising temperature on insect herbivores. Global Change Biology, 8, 1-16.

Barber NA, Marquis RJ (2011) Light environment and the impacts of foliage quality on herbivorous insect attack and bird predation. Oecologia, 166, 401-409.

Bates D, Maechler M, Bolker B, Walker S (2015) Fitting linear mixed-effects models using lme4. Journal of Statistical Software, 67(1), 1-48.

Belsky AJ (1986) Does herbivory benefit plants? A review of the evidence. American Naturalist, 127, 870-892.

Bertheau C, Brockerhoff EG, Roux-Morabito G, Lieutier F, Jactel H (2010) Novel insect-tree associations resulting from accidental and intentional biological ‘invasions’: a meta-analysis of effects on insect fitness. Ecology Letters, 13, 506-515.

Bixenmann RJ, Coley PD, Weinhold A, Kursar TA. (2016) High herbivore pressure favors constitutive over induced defense. Ecology and Evolution, 6, 6037-6049.

Brasier CM. (1992) Oak Tree mortality in Iberia. Nature, 360, 539–539.

Bryant JP, Chapin III FS, Klein DR (1983) Carbon/nutrient balance of boreal plants in relation to vertebrate herbivory. Oikos, 40, 357–368.

Bryant JP, Clausen TP, Reichardt PB, McCarthy MC, Werener RA (1987) Effects of nitrogen fertilisation upon the secondary chemistry and nutritional value of quaking aspen (*Populus tremuloides* Michx.) leaves for the large aspen tortrix (*Choristoneura conflicta* (Walker)). Oecologia, 73, 513–517.

Bugalho MN, Lecomte X, Gonçalves M, Caldeira MC, Branco M (2011) Establishing grazing and grazing-excluded patches increases plant and invertebrate diversity in a Mediterranean oak woodland. Forest Ecology and Management, 261, 2133–2139.

Bugalho MN, Ibáñez I, Clark JS (2013) The effects of deer herbivory and forest type on tree recruitment vary with plant growth stage. Forest Ecology and Management, 308, 90–100.

Burger JC, Louda SM (1994) Indirect versus direct effects of grasses on growth of a cactus (*Opuntia fragilis*): insect herbivory versus competition. Oecologia, 99, 79-87.

Caldeira MC, Ibáñez I., Nogueira C, Bugalho MN, Lecomte X, Moreira A, Pereira JS. (2014) Direct and indirect effects of tree canopy facilitation in the recruitment of Mediterranean oaks. Journal of Applied Ecology, 51, 349-358.

Callaway RM (2007) Positive interactions and interdependence in plant communities. Springer.

Campos P, Huntsinger L, Oviedo JL, Starrs PF, Díaz M, Standiford RB, Montero G (2013) Mediterranean oak woodland working landscapes. Springer.

Carnicer J, Coll M, Ninyerola M, Pons X, Sánchez G, Peñuelas J (2011) Widespread crown condition decline, food web disruption, and amplified tree mortality with increased climate change-type drought. Proceedings of the National Academy of Sciences, 108, 1474-1478.

Chacón P, Armesto JJ. (2006) Do carbon-based defenses reduce foliar damage? Habitat-related effects on tree seedling performance in a temperate rainforest of Chiloé Island, Chile. Oecologia, 146, 555-565.

Chaneton EJ, Mazia CN, Kitzberger T (2010) Facilitation vs. apparent competition: insect herbivory alters tree seedling recruitment under nurse shrubs in a steppe-woodland ecotone. Journal of Ecology, 98, 488-497.

Coley PD, Bryant JP, Chapin FS (1985) Resource Availability and Plant antiherbivore defense. Science, 230, 895.

Connell JH (1971) On the role of natural enemies in preventing competitive exclusion in some marine animals and rain forest trees. Dynamics of population, 298-312.

Corcket E, Callaway RM, Michalet R (2003) Insect herbivory and grass competition in a calcareous grassland: results from a plant removal experiment. Acta Oecologica, 24, 139–146.

Cornelissen T, Fernandes GW, Vasconcellos-Neto J (2008) Size does matter: variation in herbivory between and within plants and the plant vigor hypothesis. Oikos, 117, 1121–1130.

Crawley MJ (1997) Plant ecology. Second edition. Blackwell Science.

Cuesta B, Villar-Salvador P, Puértolas J, Rey Benayas JM, Michalet R (2010) Facilitation of *Quercus ilex* in Mediterranean shrubland is explained by both direct and indirect interactions mediated by herbs. Journal of Ecology, 98, 687-696.

Díaz M, Pulido FJ, Møller AP (2004) Herbivore effects on developmental instability and fecundity of holm oaks. Oecologia, 139, 224-234.

Dudt JF, Shure DJ (1994) The influence of light and nutrients on foliar phenolics and insect herbivory. Ecology, 75, 86-98.

Fedriani JM, García LV, Sánchez ME, Calderón J, Ramo C (2017) Long-term impact of protected colonial birds on a jeopardized cork oak population: conservation bias leads to restoration failure. Journal of Applied Ecology, 54, 450-458

Feeny P (1970) Seasonal changes in oak leaf tannins and nutrients as a cause of spring feeding by winter moth caterpillars. Ecology, 51, 565-581.

Fine PVA, Miller ZJ, Mesones I, Irazuzta S, Appel HM, Stevens MHH, Saaksjarvi I, Schultz LC, Coley PD (2006) The growth-defense trade-off and habitat specialization by plants in Amazonian forests. Ecology, 87, S150-S162.

Folin O, Ciocalteu V (1927) On tyrosine and tryptophane determinations in proteins. Journal of Biological Chemistry, 73, 627-650.

Fornoni J (2011) Ecological and evolutionary implications of plant tolerance to herbivory. Functional Ecology, 25, 399-407.

García-Cervigoni AI, Iriondo JM, Linares JC, Olano JM (2016) Disentangling facilitation along the life cycle: Impacts of plant-plant interactions at vegetative and reproductive stages in a Mediterranean forb. Frontiers in Plant Science, 7, 1-11.

Gavinet J, Prévosto B, Fernandez C (2016) Do shrubs facilitate oak seedling establishment in Mediterranean pine forest understory? Forest Ecology and Management, 381, 289-296.

Grace JB. 2006 Structural equation modeling and natural systems. Cambridge University Press, Cambridge, UK.

Hamilton JG, Zangerl AR, DeLucia EH, Berenbaum MR (2001) The carbon–nutrient balance hypothesis: its rise and fall. Ecology Letters, 4, 86-95.

Herms DA, Mattson WJ (1992) The dilemma of plants - to grow or defend. Quarterly Review of Biology, 67, 283-335.

Janzen DH. (1970) Herbivores and the number of tree species in tropical forests. American Naturalist, 104, 501-528.

Kellner KF, Swihart RK (2017) Herbivory on planted oak seedlings across a habitat edge created by timber harvest. Plant Ecology, 218, 213-223.

Ladd BM, Facelli JM (2005) Effects of competition, resource availability and invertebrates on tree seedling establishment. Journal of Ecology, 93, 968–977.

Lamarre GPA, Mendoza I, Fine PVA, Baraloto C (2014) Leaf synchrony and insect herbivory among tropical tree habitat specialists. Plant Ecology, 215, 209-220.

Leck MA, Parker T, Simpson R (2008) Seedling Ecology and Evolution. Cambridge University Press.

Lecomte X, Fedriani JM, Caldeira MC, Clemente AS, Olmi A, Bugalho MN. 2016. Too many is too bad: Long-term net negative effects of high density ungulate populations on a dominant Mediterranean shrub. Plos One 11:e0158139.

Lefcheck JS. (2016) piecewiseSEM: Piecewise structural equation modeling in R for ecology, evolution, and systematics. Methods in Ecology and Evolution, 7(5), 573-579.

Leonardsson J, Lof M, Gotmark F (2015) Exclosures can favour natural regeneration of oak after conservation-oriented thinning in mixed forests in Sweden: A 10-year study. Forest Ecology and Management, 354, 1–9.

Marquis RJ (1984) Leaf herbivores decrease fitness of a tropical plant. Science, 226, 537-539.

Martinat PJ (1987) The role of climatic variation and weather in forest insect outbreaks. – In: Barbosa P and Schultz JC (eds), Insect outbreaks. Academic Press, pp. 241–268.

Mazza CA, Izaguirre MM, Zavala J, Scopel AL, Ballaré CL (2002) Insect perception of ambient ultraviolet-B radiation. Ecology Letters, 5, 722-726.

McNaughton SJ (1983) Compensatory Plant Growth as a Response to Herbivory. Oikos, 40, 329-336.

Nakagawa S, Schielzeth H (2013) A general and simple method for obtaining *R*^2^ from generalized linear mixed-effects models. Methods in Ecology and Evolution, 4, 133–142.

Norghauer JM, Newbery DM (2014) Herbivores differentially limit the seedling growth and sapling recruitment of two dominant rain forest trees. Oecologia, 174, 459-469.

Onoda Y, Westoby M, Adler PB, Choong AMF, Clissold FJ, Cornelissen JHC, … Yamashita N (2011) Global patterns of leaf mechanical properties. Ecology Letters, 14, 301-312.

Paracchini ML, Petersen JE, Hoogeveen Y, Bamps C, Burfield I, van Swaay C (2008) High nature value farmland in Europe. An estimate of the distribution patterns on the basis of land cover and biodiversity data. Office for Official Publications of the European Communities, Luxembourg.

Pearse IS, Hipp AL (2009) Phylogenetic and trait similarity to a native species predict herbivory on non-native oaks. Proceedings of the National Academy of Sciences, 106, 18097-18102.

Perez-Ramos IM, Maranon T (2008) Factors affecting post-dispersal seed predation in two coexisting oak species: Microhabitat, burial and exclusion of large herbivores. Forest Ecology and Management, 255, 3506-3514.

Pinheiro J, Bates D, DebRoy S, Sarkar D, R Core Team (2017). nlme: Linear and Nonlinear Mixed Effects Models. R package version 3.1–131.

Pinto CA, Henriques MO, Figueiredo JP, David JS, Abreu FG, Pereira JS, … David TS. (2011) Phenology and growth dynamics in Mediterranean evergreen oaks: Effects of environmental conditions and water relations. Forest Ecology and Management, 262, 500-508.

Plieninger T, Rolo V, Moreno G (2010) Large-scale patterns of *Quercus ilex*, *Quercus suber*, and *Quercus pyrenaica* regeneration in Central-Western Spain. Ecosystems, 13, 644-660.

Puerta-Piñero C, Gómez JM, Valladares F (2007) Irradiance and oak seedling survival and growth in a heterogeneous environment. Forest Ecology and Management, 242, 462-469.

Quinn GP, Keough MJ (2002) Experimental Design and Data Analysis for Biologists. Cambridge University Press, Cambridge.

R Core Team (2017) R: A language and environment for statistical computing. R Foundation for Statistical Computing, Vienna, Austria. URL https://www.R-project.org/.

Rhoades DF (1979) Evolution of plant chemical defense against herbivores. In GA Rosenthal & DH Janzen (Eds.). Herbivores: their interactions with secondary plant metabolites (pp. 1–55). New York: Academic Press.

Rolo V, Plieninger T, Moreno G (2013) Facilitation of holm oak recruitment through two contrasted shrubs species in Mediterranean grazed woodlands. Journal of Vegetation Science, 24, 344–355.

Sampaio T, Branco M, Guichoux E, Petit RJ, Pereira JS, Varela MC, Almeida MH (2016) Does the geography of cork oak origin influence budburst and leaf pest damage? Forest Ecology and Management, 373, 33–43.

Scheipl F, Greven S, Kuechenhoff H (2008) Size and power of tests for a zero random effect variance or polynomial regression in additive and linear mixed models. Computational Statistics & Data Analysis, 52, 3283–3299.

Shipley B. (2009) Confirmatory path analysis in a generalized multilevel context. Ecology, 90, 363–368.

Sipura M, Tahvanainen J (2000) Shading enhances the quality of willow leaves to leaf beetles – but does it matter? Oikos, 91, 550–558.

Sotomayor DA, Lortie CJ (2015) Indirect interactions in terrestrial plant communities: emerging patterns and research gaps. Ecosphere, 6, 23.

Stephens AEA, Westoby M (2015) Effects of insect attack to stems on plant survival, growth, reproduction and photosynthesis. Oikos, 124, 266–273.

Talamo A, Barchuk AH, Garibaldi LA, Trucco CE, Cardozo S, Mohr F (2015) Disentangling the effects of shrubs and herbivores on tree regeneration in a dry Chaco forest (Argentina). Oecologia, 178, 847–854.

Therneau TM, Grambsch PM (2000) Modeling survival data: Extending the Cox model. Springer, New York.

Therneau T (2015) A Package for Survival Analysis in S. version 2.38.

Therneau T., Crowson C., Atkinson E. (2016) Using time dependent covariates and time dependent coefficients in the cox model. Survival Vignettes 2016.

Tiberi R, Branco M, Bracalini M, Croci F, Panzavolta T (2016) Cork oak pests: a review of insect damage and management. Annals of Forest Science, 73, 219–232.

Valladares F, Laanisto L, Niinemets U, Zavala MA (2016) Shedding light on shade: ecological perspectives of understorey plant life. Plant Ecology & Diversity, 9, 237–251.

Warton DI, Hui FKC. (2011) The arcsine is asinine: the analysis of proportions in ecology. Ecology, 92, 3–10.

White T CR. (1969) An index to measure weather-induced stress of trees associated with outbreaks of psyllids in Australia. Ecology 50, 905–909.

White TCR. (1974) A hypothesis to explain outbreaks of looper caterpillars, with special reference to populations of *Selidosema suavis* in plantations of *Pinus radiata* in New Zealand. Oecologia 16, 279–301.

White TCR. (1984) The abundance of invertebrate herbivores in relation to the availability of nitrogen in stressed food plants. Oecologia 63, 90–05.

White, TCR. (2009) Plant vigour versus plant stress: a false dichotomy. Oikos, 118, 807–808.

WRB. (2006) World Reference Base for Soil Resources. A Framework for International Classification, Correlation and Communication. World Soil Reports, 103, FAO, Rome.

Zuur AF, Ieno EN, Walker N, Saveliev AA, Smith GM (2009) Mixed effects models and extensions in ecology with R. Springer, New York.

